# Chronic Seizures Induce Sex-Specific Cognitive Deficits with Loss of Presenilin 2 Function

**DOI:** 10.1101/2021.12.16.472840

**Authors:** Kevin M. Knox, Megan Beckman, Carole L. Smith, Suman Jayadev, Melissa Barker-Haliski

## Abstract

Patients with early-onset Alzheimer’s disease (EOAD) are at elevated risk for seizures, including patients with presenilin 2 (PSEN2) variants. Like people with epilepsy, uncontrolled seizures may worsen cognitive function in AD. While the relationship between seizures and amyloid beta accumulation has been more thoroughly investigated, the role of other drivers of seizure susceptibility in EOAD remain relatively understudied. We therefore sought to define the impact of loss of normal *PSEN2* function and chronic seizures on cognitive function in the aged brain. Male and female PSEN2 KO and age- and sex-matched wild-type (WT) mice were sham or corneal kindled beginning at 6-months-old. Kindled and sham-kindled mice were then challenged up to 6 weeks later in a battery of cognitive tests: non-habituated open field (OF), T-maze spontaneous alternation (TM), and Barnes maze (BM), followed by immunohistochemistry for markers of neuroinflammation and neuroplasticity. PSEN2 KO mice required significantly more stimulations to kindle (males: p<0.02; females: p<0.02) versus WT. Across a range of behavioral tests, the cognitive performance of kindled female PSEN2 KO mice was most significantly impaired versus age-matched WT females. Male BM performance was generally worsened by seizures (p=0.038), but loss of PSEN2 function did not itself worsen cognitive performance. Conversely, kindled PSEN2 KO females made the most BM errors (p=0.007). Chronic seizures also significantly altered expression of hippocampal neuroinflammation and neuroplasticity markers in a sex-specific manner. Chronic seizures may thus significantly worsen hippocampus-dependent cognitive deficits in aged female, but not male, PSEN2 KO mice. Our work suggests that untreated focal seizures may worsen cognitive burden with loss of normal PSEN2 function in a sex-related manner.

## Introduction

Alzheimer’s disease (AD) is a devastating neurological condition that is irreversible, unstoppable, and ultimately fatal. Most cases first become symptomatic in old age (65+), but early-onset AD (EOAD) occurs in individuals over 50-years-old with an annual incidence rate of 6.3/100,000 [1]. People with EOAD typically experience worse severity and more rapid disease progression. These people experience 87-times higher risk of seizures versus same-aged people without AD [2, 3]. Although not widely recognized as a comorbidity of AD, seizures may accelerate functional decline in patients and dramatically increase caregiver burden [4]. Seizures in AD are often unprovoked focal impaired awareness seizures, meaning they have no obvious trigger or clinical signs, and therefore can go unrecognized [5]. Failure to treat seizures may additively compound neuropathology in AD. This is especially tragic because antiseizure medications (ASMs) may improve cognitive function in older people with mild cognitive impairment [6], and seizures in elderly patients with epilepsy are generally well-controlled [7], suggesting that rational selection of ASMs could be an untapped strategy to slow seizure-related functional decline in EOAD. However, few studies have defined the direct functional impacts of evoked chronic seizures in AD-related genotypes to spur intervention trials. As a result, clinical studies for ASMs in AD patients have been largely unsuccessful [8].

EOAD is associated with variants in the homologous presenilin 1 (*PSEN1*) and presenilin 2 (*PSEN2*) genes [9]. The mechanisms by which *PSEN* gene variants cause AD remains unclear [10-12], but it is known that AD patients with *PSEN2* variants also exhibit a high incidence of seizures [13, 14]. Intramembrane presenilin proteins (PS) perform several diverse biological functions within the CNS that may be relevant to seizure risk. PS proteins comprise the catalytic component of γ-secretase, which promotes the cleavage of amyloid-beta (Aβ) oligomers from the amyloid precursor protein (APP; [15]). PS proteins are localized to the endoplasmic reticulum membrane to regulate calcium (Ca^2+^) leak, a function that is independent of γ-secretase activity [16]. Loss of normal PS signaling induces defective Ca^2+^ signaling. PS/γ-secretase activity can modulate the cleavage of voltage-gated K^+^ channel β-subunits *KCNE1* and *KCNE2* [17], pointing to a potential role in modulating voltage-dependent currents. Notably, PS2 protein mediates signaling pathways distinct from PS1 [18]. Variants in both *PSEN1* and *PSEN2* lead to a biochemical loss-of-normal function [19] such that *PSEN* knockout models are relevant to *a priori* study the functional consequences of PSEN dysfunction across a lifespan. PSEN1 KO mice are however nonviable, whereas PSEN2 KO mice demonstrate normal development and are viable over 12-months-old [9]. Therefore, PSEN2 KO mice are useful to understand how global impairment of PSEN signaling influences hyperexcitability and pathology during both neurological development and aging.

PSENs are also important to assess immune-mediated contributions to AD, rather than amyloid β (Aβ) accumulation *per se*. PSEN2 is an underexplored molecular contributor to neuroinflammation due to the conserved regulation between PSEN2 and NFκB and molecules of the Toll-like receptor signaling pathway [20], both of which are heavily implicated in the inflammatory response in both epilepsy [21-23] and AD [24]. Neuroinflammation may also worsen cognitive decline in both epilepsy and AD. For example, elevated levels of IL-1b may underlie cognitive worsening, especially on hippocampus-mediated tasks [25]. IL-1b overexpression can disrupt long-term contextual and spatial memory but does not affect short-term and non-hippocampal memory [26], potentially through suppression of brain-derived neurotrophic factor (BDNF). BDNF itself indirectly regulates AMPA-type glutamate receptor localization to activated neuronal synapses required for learning and memory [27-29]. BDNF expression correlates to behavioral performance [30]. However, limited studies in AD models have defined the additive impact of chronic seizures on cognitive performance and neuropathology [31]. No study has yet demonstrated how loss of normal PSEN2 function, which primes an inflammatory response [32, 33], may impact cognitive function with chronic seizures.

The amyloid hypothesis has been the predominate framework in AD research and the existence of Aβ plaques and neurofibrillary tangles is likely fundamental to AD onset, but this hypothesis alone is insufficient to wholly reproduce all clinical features. Thus far, studies of seizures in AD models have primarily focused on the impacts of APP and tau overexpression, leaving little insight into alternative drivers of AD pathogenesis. Moreover, these studies have largely relied on the passive recording of spontaneous electrographic discharges in chronically implanted animals, which is less conducive to pair with precisely timed cognitively demanding behavioral assessments. We have previously applied the corneal kindled mouse (CKM) model of chronic focal seizures to define the dose-related anticonvulsant efficacy of ASMs in young versus aged PSEN2 KO mice [34]. This study thus sought to characterize risk for cognitive deficit in the absence of Aβ in mature adult mice to understand whether seizures may also be a causative driver cognitive decline in AD. Importantly, this investigator-controlled model of chronic focal seizures provides a valuable framework in which to directly assess functional impacts of chronic seizures with AD-associated genotypes [8]. Therefore, this study sought to define how hippocampally-mediated behavioral performance and neuropathology of adult PSEN2 KO mice (>6 months old) would be directly impacted by chronic, untreated focal seizures to establish the proof-of-principle use of CKM for behavioral testing with relevant AD-associated genetic mouse models.

## Methods

### Animals

Male and female PSEN2 KO mice were bred at the University of Washington (UW) from stock originally acquired from the Jackson Laboratory (stock #005617). All breeding at UW was between PCR-confirmed PSEN2 KO pairs. Age-matched male and female C57Bl/6J WT (stock #000664) were acquired from a similarly established breeding colony at UW, which was matched in housing conditions with the PSEN2 KO mice. This study was not testing the effects of sex hormones on behavior or protein expression; thus, we did not track estrous cycles of females or the dominant/subordinate status of males. Mice were housed on a 14:10 light cycle (on at 6 h00; off at 20 h00) in ventilated cages in corncob bedding in a manner consistent with the *Guide for the Care and Use of Laboratory Animals*. All animal work was approved by the UW Institutional Animal Care and Use Committee (protocol 4387-01) and conformed to ARRIVE guidelines [35]. Animals were provided free access to irradiated chow (PicoLab Rodent Diet 20-5053), filtered water, and enrichment, except during periods of behavioral manipulation. All testing was performed between 7h00 and 19h00.

### Corneal Kindling

Two groups of 6-month-old mice from each genotype (PSEN2 KO and WT) underwent corneal kindling. Kindling group sizes are presented in Table 1. Tetracaine-HCl (0.5%) analgesic was applied bilaterally to the corneas immediately preceding a 3 s, 60 Hz sinusoidal electrical stimulation. Behavioral manipulation was identical for kindled and sham groups, except no electrical stimulation was delivered to sham mice. Stimulation intensity was based on the subconvulsive minimal clonic seizure thresholds for age-, sex- and genotype-matched mice [34]. Males were stimulated at 3.0 mA; females were stimulated at 2.6 mA. Kindled seizure severity was scored according to a modified Racine scale [36, 37]. Mice were kindled until they achieved criterion, defined as five consecutive stage 5 seizures [34]. The total number of mice that attained kindling criterion for each group are presented in Table 1. Mice that did not reach criterion were excluded from further study. Following corneal or sham kindling, mice were divided into two testing cohorts (Table 1). Cohort 1 completed the open field (OF) test and T-maze continuous alternation task (TM). Cohort 2 completed the Barnes maze (BM).

**Table 1.** Experimental group distribution and group sizes for 6-month-old WT and PSEN2 KO mice studies.

| **Experimental Group and Sex** | **Wild-Type – Sham Kindled** | **Wild-Type - Kindled** | **PSEN2 KO – Sham-Kindled** | **PSEN2 KO - Kindled** |
| --- | --- | --- | --- | --- |
| Male (all; WT n = 55; KO = 51) | 23 | 32 | 19 | 32 |
| Female (all; WT n = 41; KO = 53) | 18 | 23 | 21 | 32 |
| Males Reaching Kindling Criterion (% of total) | 0 (0%) | 25 (78.1%) | 0 (0%) | 24 (75%) |
| Females Reaching Kindling Criterion (% of total) | 0 (0%) | 21 (91.3%) | 0 (0%) | 22 (68.8%) |
| Cohort 1 Males for OF/TM | 10 | 12 | 8 | 12 |
| Cohort 1 Females for OF/TM | 9 | 12 | 9 | 9 |
| Cohort 2 Males for BM | 13 | 13 | 11 | 12 |
| Cohort 2 Females for BM | 9 | 9 | 12 | 13 |

A separate cohort of old WT (n=14) and PSEN2 KO (n=11) female mice (>14-months-old) were enrolled for corneal kindling to determine whether advanced age would be associated with sustained slower kindling rate. Aged females were similarly kindled as for 6-month-old mice. These older females were not assessed for behavioral performance or immunohistochemical analysis but were only included to define the interaction between advanced age and PSEN2 KO on kindling susceptibility.

### Seizure Duration

The kindled seizure duration was recorded from fully kindled mice of Cohort 1 following the 17-day stimulation-free period in which OF and TM tasks were completed to confirm the stability of the fully kindled state. The duration of seizure-induced behavioral arrest for mice that presented with a secondarily generalized (Racine stage 4 or 5) seizure at this time point post-kindling was recorded by an investigator blinded to genotype and kindling status [34].

### Open Field (OF)

In this task, each mouse was placed into an OF Plexiglas chamber (40L × 40W × 30H cm) equipped with infrared sensors to detect animal movement for 10 min, including the total distance travelled (cm), vertical activity counts, and horizontal activity counts in both the center and perimeter [38-42].

### T-maze (TM)

The TM continuous alternation task is a land-based task well-suited to assess hippocampally-mediated spatial working memory of mice [43]. Each TM test consisted of 6-8 trials. Mice were allowed to alternate spontaneously between the left and right goal arms (30 × 10 cm) of a T-shaped epoxy-coated wooden maze (start alley 30 × 10 cm) throughout a 15-trial session. On the testing day, each mouse was contained in a starting box (10 × 8 cm) at the entry to the start arm. At the start of each trial, a partition door was opened to allow the mouse to move down the starting arm and choose between 2 goal arms, left or right. Once a mouse entered a particular goal arm, a door was lowered quietly to block further exploration of the opposite arm. Arm choices were recorded manually by a non-interfering observer. After 30 seconds, the mouse was returned to the starting box and the start alley and goal arms cleaned with 30% isopropyl alcohol and dried to remove any olfactory cues. The mouse was then rechallenged to explore the maze again, and the arm selection recorded. The spontaneous alternation rate was calculated as the ratio between the alternating choices and total number of choices during all trials, with 60% considered successful TM performance [44].

### Barnes Maze (BM)

The BM is a rodent test of spatial learning and memory [45, 46] that is particularly well suited for rodents with epilepsy [47]. The platform consisted of a 68.6 cm diameter, white Plexiglas circular platform with twenty equally spaced escape holes and a hidden, black escape chamber attached below one of the twenty holes. Extra-maze spatial cues remained in a constant location, including the investigator. Bright lights (∼1200 lux) were placed overhead of the maze as an aversive stimulus. The mildly aversive stimulus formed the basis to motivate each mouse to escape and thus learn the location of the escape chamber. The location of the escape hole remained the exact same for each individual mouse during the four acquisition days, but was randomly assigned to one of 8 coordinates (N, NW, S, SW, E, NE, W, SE) for each mouse in each experimental group. Before each trial, the BM platform, the escape chamber, and the start tube were cleaned with 30% isopropyl alcohol and air dried to remove any olfactory cues.

Testing on the BM proceeded in three phases [48]: the habituation phase (day 0), an acquisition phase (days 1-4), and a probe trial of long-term memory (day 7). The habituation day was designed to familiarize each mouse with the escape chamber to reinforce escape from the bright lights illuminating the maze. The acquisition phase was designed to teach each mouse the location of the escape chamber over a series of twelve trials. During the probe trial, the escape chamber was removed, and one trial performed to determine if the animal remembered where the target escape hole was located. The latency to first encounter with escape chamber hole and number of holes explored (errors) before first encounter with the target/escape chamber hole was manually recorded by a non-interfering observer during each acquisition and probe trial.

### Euthanasia

One hour after the conclusion of TM and BM testing, mice were euthanized by live decapitation in accordance with IACUC guidelines and brains were collected, flash frozen on dry ice, and stored at −80°C. Brains were later sectioned on a Leica CM1860 cryostat into 20-µm thick coronal slices from Bregma −1.70 to −2.10 to capture the dorsal hippocampus for subsequent immunohistochemistry. Two rostral and two caudal sections/mouse were mounted on a Superfrost slide (Fisher Scientific).

### Immunohistochemistry

After cryosectioning, slides were fixed in 4% Formal Fixx (ThermoFisher) for 10 minutes (22°C) then dehydrated in 50% ethanol for 30 min. After dehydration, the slides were washed (3 × 10 min) in 0.1 M PBS before being incubated in a 10% Goat Serum in 5% Triton-X PBS solution under coverwells in a humid chamber for 2 hours. Briefly, NeuN (1:300; MAB377X, Millipore) and GFAP (1:1000; C9205 Sigma-Aldrich) antibodies were applied under 200 mL coverwells in a 5% Goat Serum/1xPBS solution overnight at 4°C [40, 49]. The following day, coverwells were removed and slides washed in PBS (3 × 10 min) before being coverslipped with Prolong Gold with DAPI (ThermoFisher).

Slides were similarly processed for BDNF with the exception that slides were blocked with 10% sheep serum in 5% Triton-X PBS under coverwells in a humid chamber for 2 hours and slides were incubated with primary antibodies for BDNF (1:500; ab226843, Abcam) and NeuN (1:300; MAB377X, Millipore). The secondary antibody (1:1000; C2306 Sigma Aldrich) was applied in 5% sheep serum/1x PBS overnight at 4°C.

Photomicrographs were captured with a fluorescent microscope (Leica DM-4) with a 20x objective (80x final magnification). Acquisition settings were held constant throughout. NeuN, GFAP, and BDNF expression levels, given as average area percentage, were automatically measured using Leica Thunder software.

### Statistics

Percent of mice that attained kindling criterion was compared within sexes by repeat measures ANOVA. Total numbers of stimulations to attain kindling criterion and kindled seizure duration was compared between genotypes within sexes by t-test. Performance on the OF, TM, and BM were compared by two-factor ANOVA (kindling x genotype) within sexes, with *post-hoc* Tukey’s tests when applicable. Immunohistochemical assessments were also compared by two-factor ANOVA (kindling x genotype) within sexes, with *post-hoc* Tukey’s tests when applicable. All analysis was conducted with GraphPad Prism version 8.0 or later, with p<0.05 considered significant.

## Results

### Kindling acquisition was significantly delayed in 6-month-old PSEN2 KO mice

Corneal kindling of 6-month-old male and female PSEN2 KO mice, and aged PSEN2 KO females, was well-tolerated, consistent with the kindling of age-matched WT mice (Figure 1). There was a significant time x genotype interaction on kindling acquisition in both 6-month-old male and female PSEN2 KO mice (males: F_(11,682)_=2.216, p=0.0123; females: F_(6,306)_=3.384, p=0.003), demonstrating that loss of normal PSEN2 function substantially slows kindling rate (Figure 1A and Figure 1B). Kindling acquisition rate of aged PSEN2 KO females was not significantly different from WT (Figure 1C). The percentage of 6-month-old male and female PSEN2 KO mice that attained kindling criterion was also significantly different between genotypes (Figure 1D and 1E; males: F_(23, 23)_ = 32.50, p<0.0001; females: F_(23, 23)_ = 14.10, p<0.0001). The differences in percentage of fully kindled mice between genotypes were apparent by the 13^th^ (males) and 18^th^ stimulation (females). However, the percent of fully kindled aged PSEN2 KO female mice was not different from WT (Figure 1F). Both male and female PSEN2 KO mice aged 6-months required significantly more stimulations to achieve criterion (Figure 1G and 1H - males: KO 17.1±0.808 vs WT 14.6±0.689, t=2.32, p=0.0243; female KO 11.3± 0.9 vs female WT 8.85±0.5, t=2.59, p<0.02). However, this difference was not evident in aged female PSEN2 KO mice (Figure 1I). Thus, loss of normal PSEN2 function leads to resistance to corneal kindling that is lost with advanced age.

**Figure 1.**
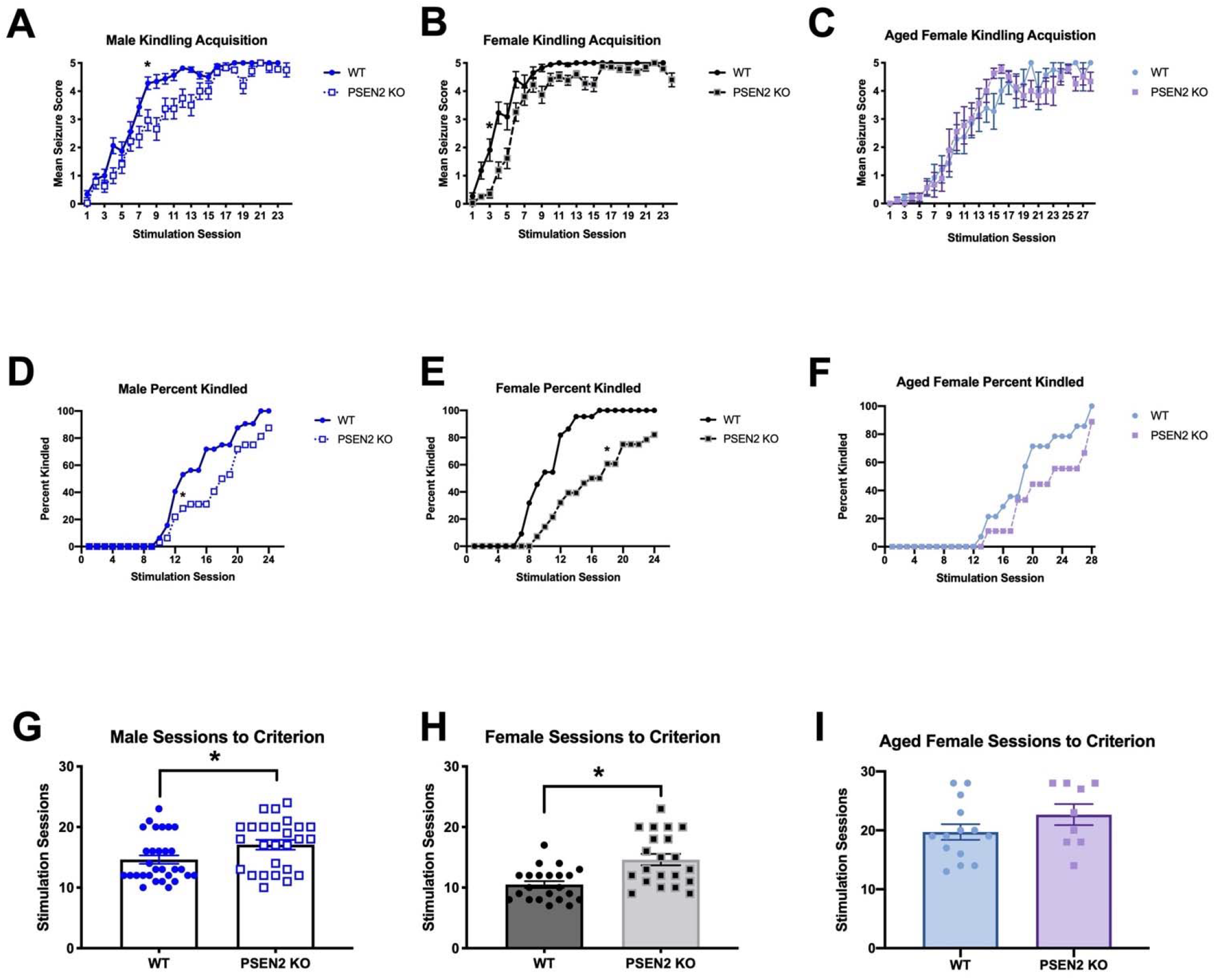
Loss of normal PSEN2 function delays corneal kindling acquisition in both male and female 6-month-old mice. Further, there is a normalization in the susceptibility to corneal kindling with advanced age in female PSEN2 KO mice. There was a significant delay in the corneal kindling acquisition rate for both A) male (F_(11,682)_=2.216, p=0.0123); and B) female (F_(6,306)_=3.384, p=0.003) PSEN2 KO mice compared to WT mice. C) However, the rate of kindling in advanced age PSEN2 KO female mice is no different from age-matched WT females (F _(1, 21)_ = 0.2294, p>0.05). The percentage of D) male (F_(23, 23)_ = 32.50, p<0.0001) and E) female (F_(23, 23)_ = 14.10, p<0.0001) PSEN2 KO mice aged 6-months-old showed a delay in reaching kindling criterion when compared to age-matched WT mice. F) However, by 14-months-old or greater, there was no longer a difference in the percentage of fully kindled female mice across time. The differences in percentage of fully kindled 6-month-old mice between genotypes were apparent by the D) 13^th^ stimulation session in males and E) 18^th^ stimulation in females. F) With advanced age, this difference in kindled percentage was no longer apparent in PSEN2 KO and WT female mice, suggesting an age-related impact of loss of normal PSEN2 function on seizure susceptibility. Finally, both male (G) and female (H) 6-month-old PSEN2 KO mice required significantly more stimulation sessions to reach kindling criterion (males: t=2.317, p=0.0243; females: t=3.797, p=0.0005). I) Conversely, advanced age female mice were not differentially susceptible to kindling, and there was no difference in the number of sessions to criterion for PSEN2 KO mice aged >14-months-old (t=1.349, p>0.05). * Indicates significantly different from WT, p<0.05.

### Open field exploration was worsened in female PSEN2 KO mice with chronic kindled seizures

The OF is useful to assess cognitive function in AD mouse models [50]. Both kindling and genotype significantly influenced the total distance traveled by male PSEN2 KO versus WT mice (Figure 2A; kindling - F_(1, 38)_=39.52, p<0.0001, genotype - F_(1, 38)_=4.682, p=0.0368), but there was no genotype x kindling interaction. *Post-hoc* tests indicated that sham-kindled PSEN2 KO and WT male mice did not significantly differ, nor did kindled PSEN2 KO and WT male mice. Like males, both kindling and genotype significantly influenced female mouse performance in the OF test (Figure 2B). There was a main effect of genotype (F_(1, 35)_=18.07, p=0.0002) and kindling (F_(1, 35)_=7.758, p=0.0086). In contrast to males, *post-hoc* tests did reveal significant differences between sham-kindled PSEN2 KO and WT female mice (p=0.035), and between kindled PSEN2 KO and WT female mice (p=0.0155). Therefore, total OF exploration was only significantly affected by genotype in female PSEN2 KO, but not male, mice.

**Figure 2.**
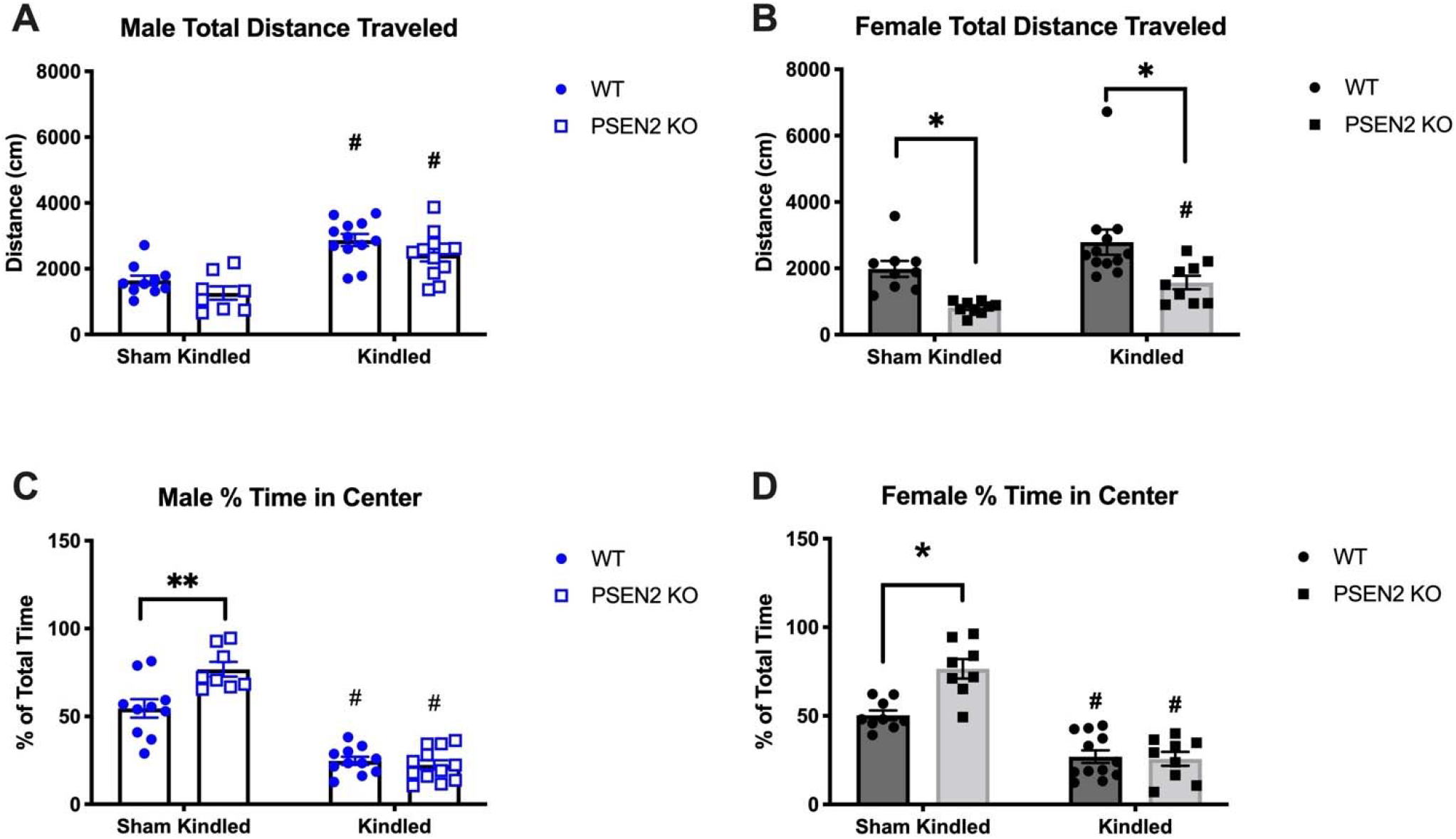
Male and female mice with loss of normal PSEN2 function (PSEN2 KO) aged over 6-months-old underwent corneal kindling or sham kindling to induce a history of chronic seizures before being challenged in the non-habituated open field (OF) test of spatial memory and anxiety-like behaviors. A) Fully kindled male PSEN2 KO and WT mice generally exhibited more total distance travelled when compared to genotype-matched sham controls (F_(1, 38)_=39.52, p<0.0001). There were no post-hoc differences between kindled PSEN2 KO and WT mice. B) There was a significant effect of kindling on total distance travelled for both female PSEN2 KO and WT mice when compared to genotype-matched sham controls (F_(1, 35)_=7.758, p=0.0086). There was also a main effect of genotype (F_(1, 35)_=18.07, p=0.0002) and *post-hoc* tests demonstrated that both sham and kindled female PSEN2 KO mice travelled less total distance when compared to kindling-matched WT controls. C) Analysis of center versus perimeter OF exploration (i.e., thigmotaxis) demonstrated a kindling x genotype interaction, with kindled male mice spending significantly less time in the center of the open field compared to sex-matched sham-kindled mice (F_(1, 37)_=11.24, p<0.002;). D) Analysis of center versus perimeter OF exploration demonstrated a kindling x genotype interaction, with kindled female mice spending significantly less time in the center of the open field compared to sex-matched sham-kindled mice (F_(1, 34)_=11.75, p<0.002). However, this change in OF exploration preference was most measured in female PSEN2 KO kindled mice. * Indicates significantly different from WT, p<0.05; # indicates significantly different from genotype-matched sham kindled group, p<0.05.

The OF is also useful to assess anxiety-like behavioral comorbidities by quantifying thigmotaxis [51], including in fully kindled mice [38]. Both male and female kindled mice demonstrated reduced total time exploring the OF center versus sham-kindled mice (Figure 2C and 2D; males: F_(1, 37)_=11.24, p<0.002; females: F_(1, 34)_=11.75, p<0.002). Kindled females generally spent less exploration time in the OF center versus sham-kindled mice (Figure 2D). Interestingly, sham-kindled female PSEN2 KO mice spent more time in the OF center compared to WT sham-kindled mice (p=0.0005) yet kindling caused female PSEN2 KO mice to spend the same amount of their total time in the OF center versus WT mice (p>0.9). Given that sham-kindled PSEN2 KO mice spent more total time in the OF center, the kindling-induced reduction in OF center zone exploration was most measured in PSEN2 KO females, suggesting that kindling more significantly worsened anxiety-like behavior of female PSEN2 KO mice.

### T-maze continuous alternation task performance was only worsened in female PSEN2 KO female mice with chronic kindled seizures

Performance of sham-versus kindled-mice on the land-based TM continuous alternation task was compared within sexes (Figure 3A and 3B). The number of TM response errors did not significantly differ between genotypes (F_(1, 40)_ = 0.4808, p>0.5) or kindling status (F_(1, 40)_ = 0.2305, p>0.6) in male mice (Figure 3A). Conversely, female mice demonstrated a significant main effect of kindling (F_(1, 36)_ = 4.482, p<0.05) and a genotype x kindling interaction on TM performance (F_(1, 36)_=5.52, p=0.024, Figure 3B). *Post-hoc* analysis revealed that fully kindled PSEN2 KO female mice performed worse than sham kindled PSEN2 KO female mice (p=0.0221) and there was a non-significant trend for worsening of performance relative to WT kindled female mice (p=0.086). Thus, TM performance was only worsened with chronic seizures in female PSEN2 KO mice.

**Figure 3.**
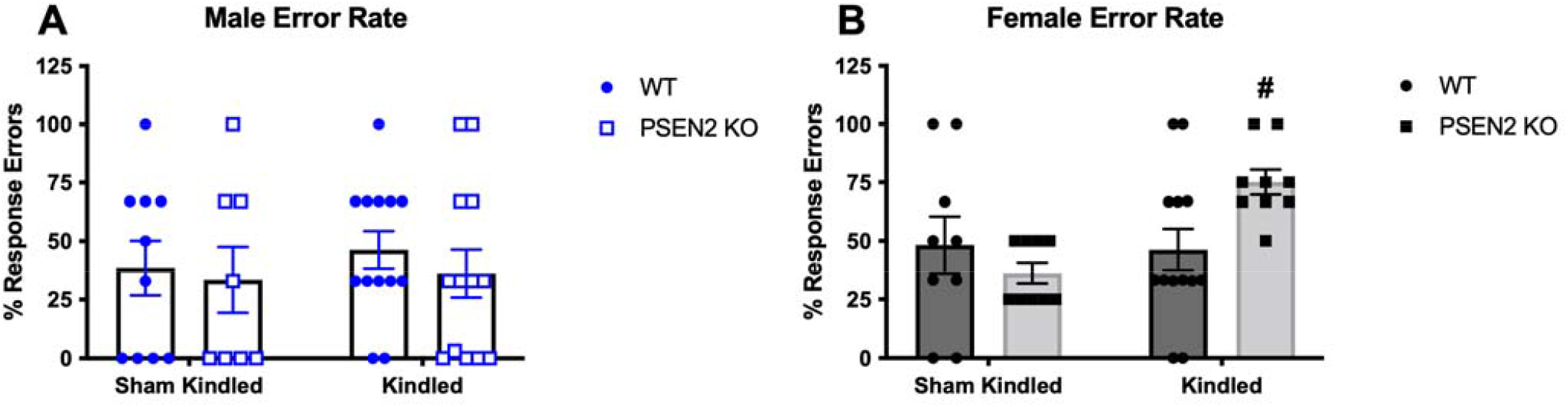
Male and female mice with loss of normal PSEN2 function (PSEN2 KO) aged over 6-months-old underwent corneal kindling or sham kindling to induce a history of chronic seizures before being challenged in the T-maze continuous alternation test of spatial working memory. A) Chronic seizures do not worsen the performance of PSEN2 KO male mice in the T-maze continuous alternation task. There was no significant genotype x kindling interaction in the male mice (F_(1, 40)_ = 0.054, p>0.8). B) Conversely, there was a significant genotype x kindling interaction in female mice (F_(1, 36)_=5.52, p=0.024). Female kindled PSEN2 KO mice made significantly more errors than the age-matched sham-kindled female PSEN2 KO mice (# indicates p<0.05).

### Behavioral seizure duration was not different between fully kindled PSEN2 KO versus WT mice

Mice in Cohort 1 had a 17-day stimulation-free period during the OF and TM test period. The duration and stability of fully kindled evoked behavioral seizures was thus confirmed after completing behavioral studies [34]. The fully kindled seizure duration was not significantly different between genotypes in either sex (not shown): male PSEN2 KO (mean±SEM – 37.2±2.43 sec) vs age-matched WT mice (38.1±3.13 sec; t=0.235, p>0.8); female PSEN2 KO (45.6±3.89 sec) versus age-matched WT mice (38.6±2.48 sec; t=1.57, p>0.1). Thus, evoked seizures were both stably induced and no different in duration between genotypes up to 17 days post-kindling, confirming that kindling-induced network hyperexcitability induced by kindling itself was not lost during the behavioral testing period.

### Barnes maze performance was worsened by chronic kindled seizures in PSEN2 KO female mice

The fully kindled (or sham kindled) mice of Cohort 2 were used to assess performance on the BM 4-6 weeks after corneal kindling. Performance on the BM for each sex was assessed by two measures over the four trial days and final probe trial on Day 7 (Figure 4): latency to first encounter with the escape hole and the number of holes explored to locate the escape chamber (errors). Neither kindling nor genotype significantly affected the latency to first encounter with the escape platform of male WT and PSEN2 KO mice (Figure 4A). Males all significantly improved their latency to first encounter on the BM with time (F_(4,176)_=21.59, p<0.0001). However, kindling generally increased the number of erroneously explored escape holes in males, regardless of genotype (Figure 4C; F_(3, 44)_=7.15, p=0.0005). There was no kindling group x number of trial day interaction for males on the BM (F_(12,176)_=0.56, p>0.8). Conversely, in the female PSEN2 KO and WT mice, chronic seizures markedly impacted BM performance in a genotype-specific manner. First, there was an interaction between kindling and trial days on the latency to first encounter with the correct escape hole (Figure 4B; F_(12,148)_=2.142, p=0.0175). Sham-kindled WT female mice were significantly slower to initially locate the hidden platform relative to kindled WT female mice (p=0.0001) and kindled PSEN2 KO mice were slower than kindled WT female mice (p=0.0138), but sham KOs were not different from kindled KOs. Second, the number of erroneously explored escape holes was increased with kindling history (Figure 4D; F_(3,37)_=12.92, p<0.0001). There were significant differences in the number of erroneously explored escape holes between kindled WT and kindled PSEN2 KO female mice on the first trial day (p=0.001) and between sham-kindled WT and kindled WT female mice on the probe trial day (Day 7, p=0.0107). Thus, chronic kindled seizures only negatively affected BM performance of PSEN2 KO females; kindled male PSEN2 KO mice were unaffected.

**Figure 4.**
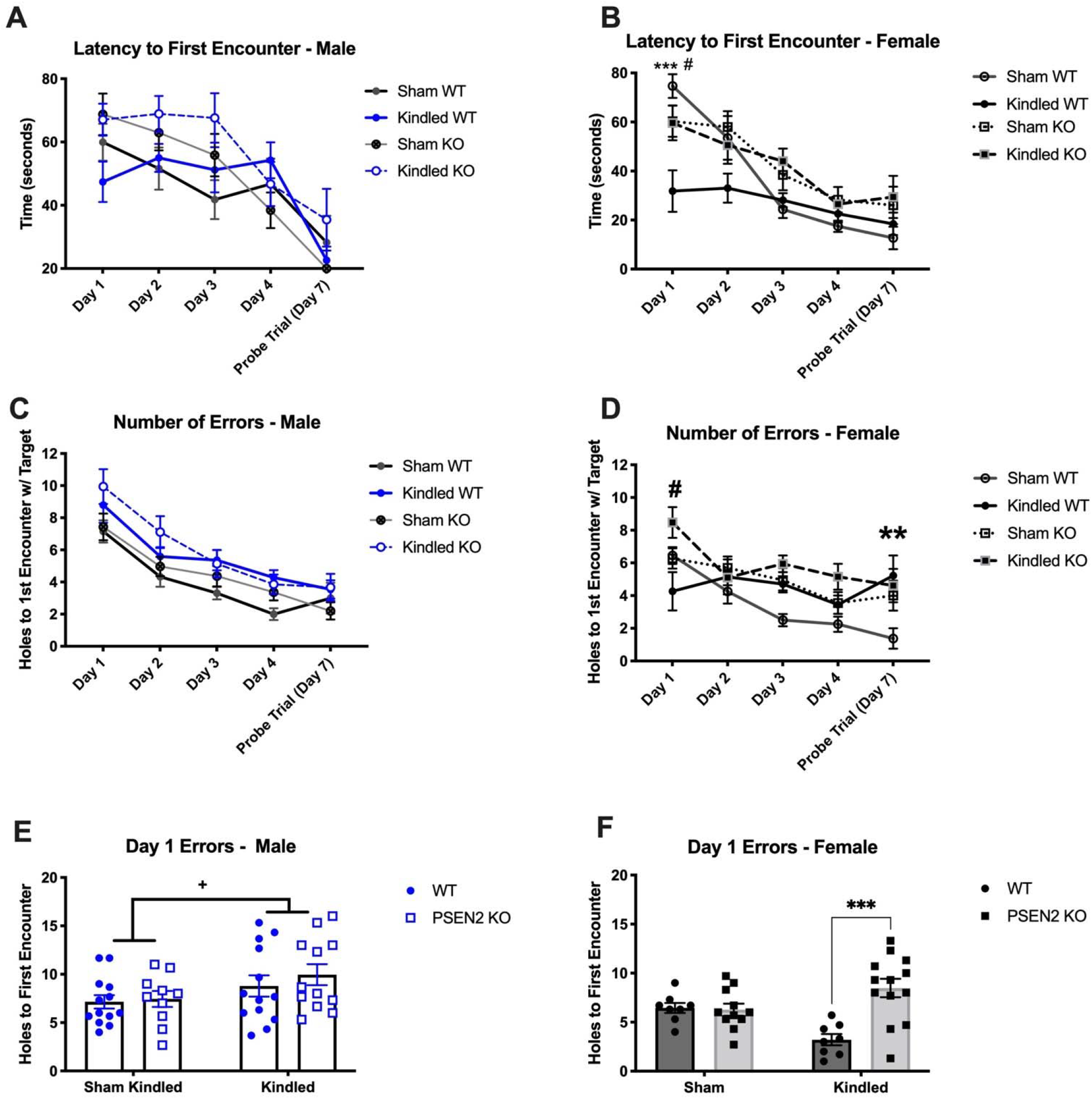
Male and female mice with loss of normal PSEN2 function (PSEN2 KO) aged over 6-months-old underwent corneal kindling or sham kindling to induce a history of chronic seizures before being challenged in the land-based, hippocampally-mediated Barnes maze (BM) test of learning and memory. BM performance was only worsened by chronic kindled seizures in PSEN2 KO female mice. A) Male mice generally demonstrated a time-related reduction in latency to first encounter with the escape chamber (F_(4,176)_=21.59, p<0.0001). However, the number of erroneously explored escape holes was generally worsened in males by kindling (Figure 4C; F_(3, 44)_=7.15, p=0.0005) but there was no interaction between the kindling groups and number of days on the BM (F_(12,176)_=0.56, p>0.8). B) Female mice generally demonstrated a time-related reduction in latency to first encounter with the escape chamber (F_(12,148)_=2.142, p=0.0175). *Post-hoc* test demonstrated that sham-kindled WT female mice were significantly slower to initially locate the hidden platform relative to kindled WT female mice (p=0.0001) and kindled PSEN2 KO mice were slower than kindled WT female mice (p=0.0138), but sham KOs were not different from kindled KOs. D) There was also a significant main effect of kindling in female mice on the number of erroneously explored escape holes (Figure 4D; F_(3,37)_=12.92, p<0.0001). E) Kindling was generally associated with worsened performance of male mice on the first testing day, regardless of genotype (F_(1,44)_=4.676, p=0.0361) but there was no effect of genotype F_(1, 44)_ = 0.554, p>0.4). F) On the converse, there was a significant kindling x genotype interaction in female mice (F_(1, 36)_ = 11.91, p=0.0014), with kindled PSEN2 KO females committing more errors on the first trial day relative to kindled WT females (p=0.0002). * Indicates significantly different from WT mice, p<0.05; # indicates significantly different from genotype-matched mice, p<0.05; + indicates significant main effect of kindling, p<0.05.

Kindled seizures negatively affected short-term memory of 6-month-old mice in a sex-specific manner, as measured by the day 1 trial performance of male and female mice (Figure 4E and 4F). Specifically, kindling generally worsened performance of male mice on the first testing day, regardless of genotype (F_(1,44)_=4.676, p=0.0361). Conversely in female mice, there was a significant kindling x genotype interaction (Figure 4F; F_(1, 36)_ = 11.91, p=0.0014); kindled PSEN2 KO females committed more errors on the first trial day compared to kindled WT females (p=0.0002). Thus, a history of chronic kindled seizures most substantially worsened the Day 1 performance of PSEN2 KO female mice aged over 6-months old, further demonstrating sex-specific functional impacts of chronic kindled seizures in PSEN2 KO mice.

### BDNF expression is reduced with chronic kindled seizures in PSEN2 KO mice

Kindled female, but not male, PSEN2 KO mice were most significantly impaired on the BM and TM tasks, suggesting that hippocampally-mediated memory deficits are specifically worsened by chronic seizures and loss of normal PSEN2 function only in female mice. We thus sought to define the extent to which the molecular marker of synaptic plasticity (BDNF) was impacted by chronic kindled seizures in WT and PSEN2 KO mice (Figure 5). BDNF expression was impacted by kindling and genotype in male mice in areas CA1 (Figure 5A; F_(1,72)_=4.087, p=0.0469) and CA3 of dorsal hippocampus (Figure 5B; (F_(1,72)_=4.504, p=0.037). BDNF expression was only significantly different in area CA1 between sham and kindled WT male mice (p=0.0348) and between kindled WT and kindled PSEN2 KO mice (p=0.0272), suggesting that chronic kindled seizures do not upregulate BDNF expression in PSEN2 KO mice. BDNF expression in the DG of kindled and sham kindled male mice of either genotype was no different.

**Figure 5.**
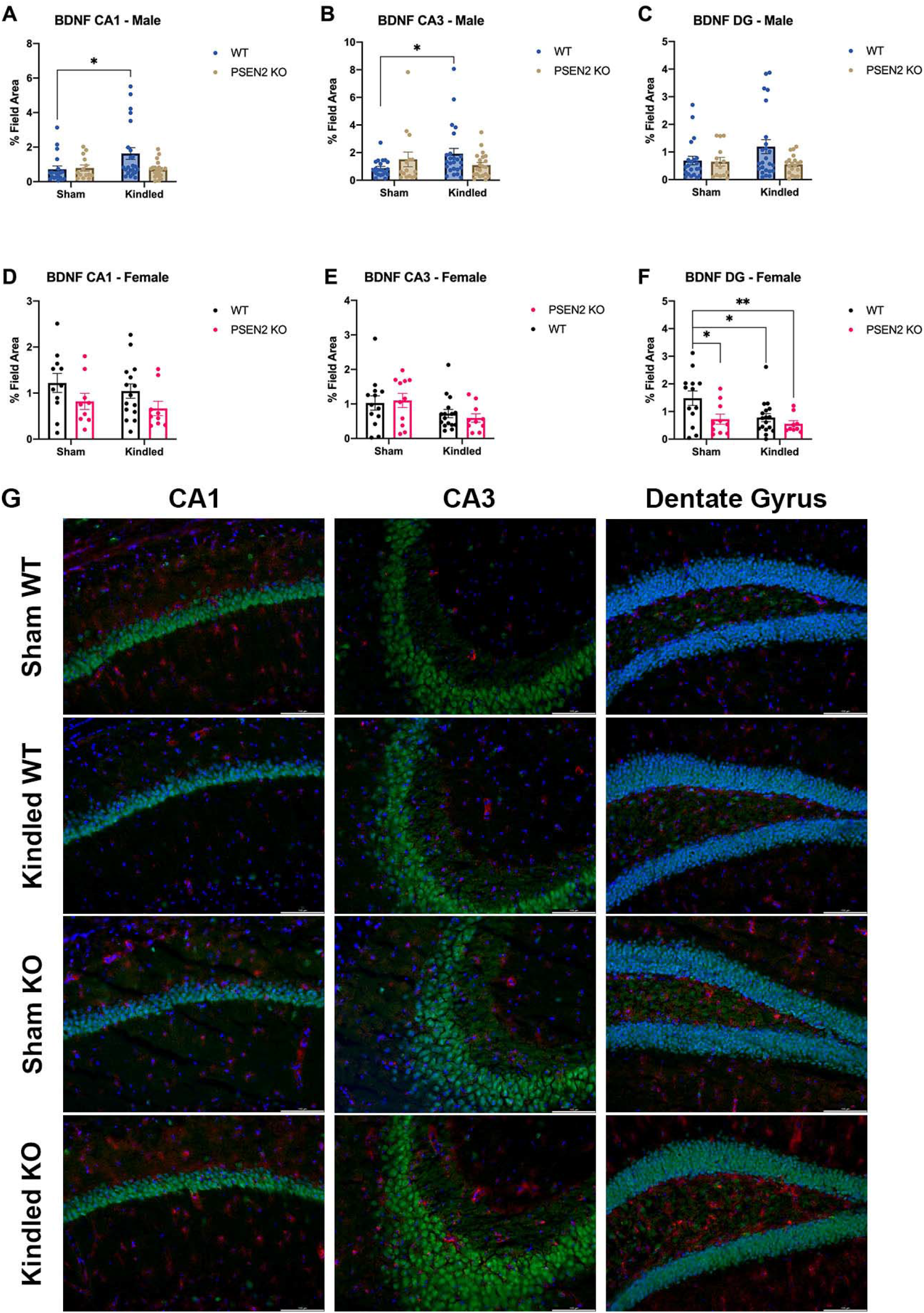
Male and female mice with loss of normal PSEN2 function (PSEN2 KO) aged over 6-months-old underwent corneal kindling or sham kindling to induce a history of chronic seizures before being challenged in a BM learning and memory test and euthanized 1 hour after the final testing day. The expression of brain derived neurotrophic factor (BDNF) was then assessed by immunohistochemistry in dorsal hippocampal structures (CA1, CA3, and Dentate Gyrus (DG)). A) There was a significant kindling x genotype effect on BDNF expression in male mice in area CA1 (F_(1,72)_=4.087, p=0.0469). *Post-hoc* tests revealed significant differences in BDNF expression between sham and kindled WT male mice (p=0.0348) and between kindled WT and kindled PSEN2 KO mice (p=0.0272). B) There was a significant kindling x genotype effect on BDNF expression in male mice in area CA3 of dorsal hippocampus (F_(1,72)_=4.504, p=0.037). C) There were no significant differences in BDNF expression in the DG of kindled and sham kindled male mice of either genotype. D) There was a significant kindling x genotype effect on BDNF expression in female mice in area CA1 (F_(1,46)_=4.607, p=0.038). E) There was a significant main effect of kindling on BDNF expression in female mice in area CA3 (F_(1,46)_=5.716, p=0.021). F) Both kindling and PSEN2 KO genotype influenced BDNF expression in DG (genotype: F_(1,46)_=6.068, p=0.0177; kindling: F_(1,46)_=4.734, p=0.0349), but there was no significant interaction between factors. Further, *post-hoc* tests only revealed significant differences in DG between sham kindled and kindled WT female mice (p=0.0358). G) Representative photomicrographs of immunohistochemical detection of BDNF in mouse brain in WT and PSEN2 KO sham- and fully kindled mice. * Indicates significantly different, p<0.05.

In female mice, PSEN2 genotype significantly influenced BDNF expression in area CA1 (Figure 5D, F_(1,46)_=4.607, p=0.038), whereas kindling significantly influenced this expression in CA3 (Figure 5E, F_(1,46)_=5.716, p=0.021). Lastly, both kindling history and genotype independently affected BDNF expression in DG (Figure 5F, genotype: F_(1,46)_=6.068, p=0.0177; kindling: F_(1,46)_=4.734, p=0.0349), but there was no significant interaction between factors. Further, *post-hoc* tests only revealed significant differences in DG between sham kindled and kindled WT female mice (p=0.0358), as well as sham kindled WT mice versus kindled PSEN2 KO female mice. While WT male mice demonstrated seizure-induced increases in BDNF expression in area CA1, this was not similarly observed in female WT mice with kindled seizures. Notably, female PSEN2 KO mice did not show kindling-induced reductions in DG BDNF expression as observed in WT female mice, further suggesting that chronic kindled seizures exert sex-specific differential effects on synaptic plasticity-associated protein expression with loss of normal PSEN2 expression.

### Fully kindled PSEN2 KO female mice demonstrate blunted seizure-induced astrogliosis

Chronic kindled seizures induce reactive astrogliosis in WT male mice [49]. Thus, we sought to quantify the degree of kindling-induced reactive astrogliosis within dorsal hippocampus of male and female WT and PSEN2 KO mice immediately after Day 7 BM testing (Figure 6A-K). Kindling-induced hippocampal astrogliosis in aged male mice was detected within both genotypes (Figure 6A and 6B), with a significant effect of kindling in CA1 (F _(1, 67)_ = 20.65, p<0.0001) and DG (F _(1, 68)_ = 32.23, p<0.0001). There were no significant differences in CA3 for any male or female group assessed. Kindling induced significant astrogliosis in both PSEN2 KO and WT male mice over 6-months-old. Further, there were no significant changes in NeuN immunoreactivity in male PSEN2 KO or WT kindled mice versus sham in either analyzed brain region (Figure 6C and 6D), suggesting that kindling of PSEN2 KO mice did not induce neurodegeneration, similar to our prior findings in young-adult WT male mice [49].

**Figure 6.**
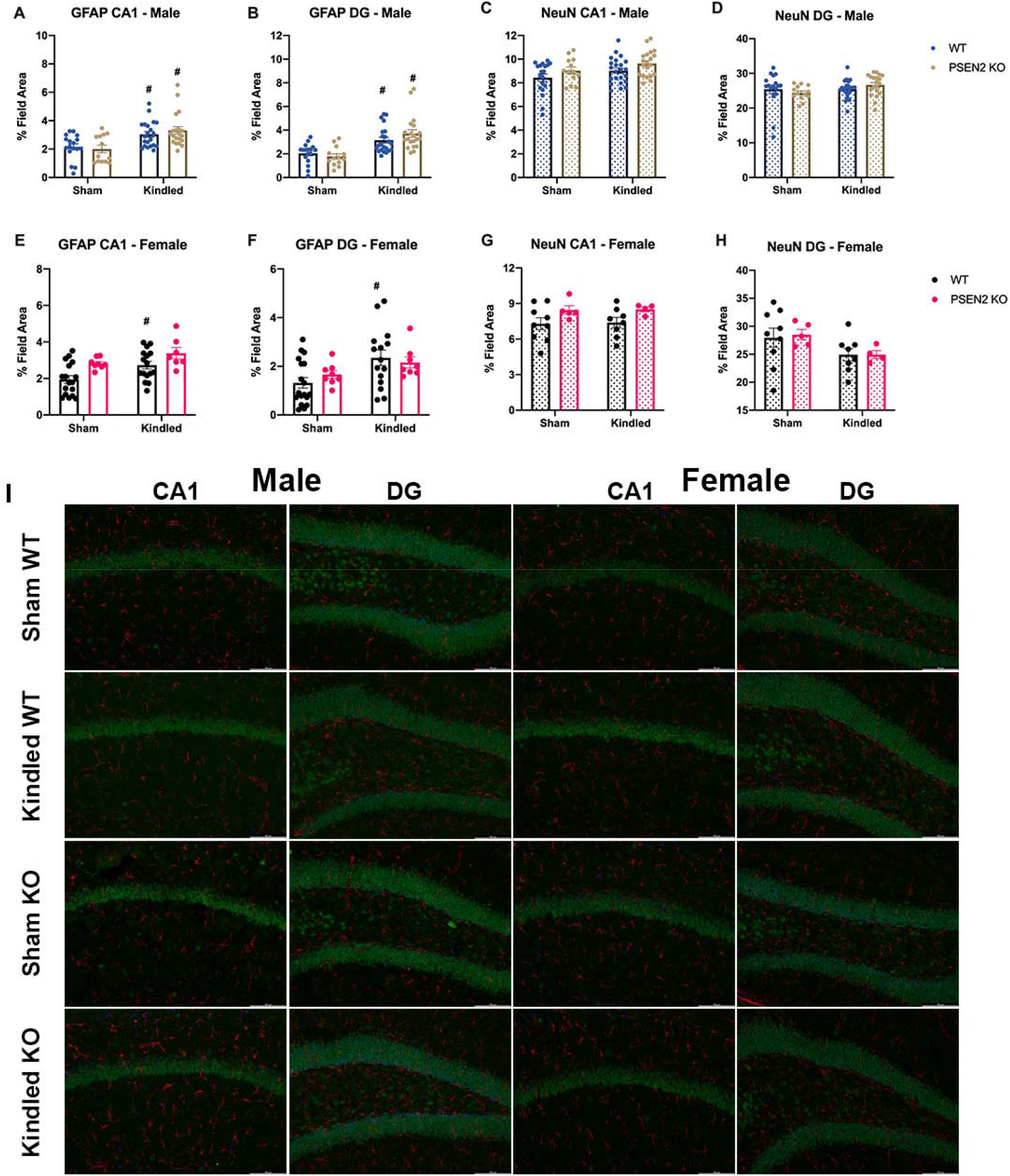
Male and female mice with loss of normal PSEN2 function (PSEN2 KO) aged over 6-months-old underwent corneal kindling or sham kindling to induce a history of chronic seizures before being challenged in a BM learning and memory test and euthanized 1 hour after the final testing day. The extent of reactive gliosis (as measured by GFAP, A-B; E-F) and neuronal density (as measured by NeuN; C-D: G-H) were evaluated in dorsal hippocampal structures (CA1 and Dentate Gyrus (DG)). There were no significant differences in area CA3, thus this region is not presented. A) There were significant kindling-induced increases in GFAP immunoreactivity in CA1 in both WT and PSEN2-KO mice (F _(1, 67)_ = 20.65, p<0.0001). B) There were significant kindling-induced increases in GFAP immunoreactivity in DG in both WT and PSEN2-KO mice (F _(1, 68)_ = 32.23, p<0.0001). C and D) There were no significant differences in NeuN immunoreactivity in CA1 and DG in male mice. E) There were significant kindling-induced increases in GFAP immunoreactivity in CA1 in both WT and PSEN2-KO female mice (F_(1, 44)_ = 7.576, p=0.0086), as well as a main effect of genotype (F_(1, 44)_ = 9.868, p=0.003) on GFAP immunoreactivity in area CA1. *Post-hoc* analysis demonstrated stimulation-induced increases in GFAP immunoreactivity in WT female mice only (p=0.030). F) There was only a main effect of genotype on DG astrogliosis (F _(1, 45)_ = 7.057, p=0.0109; Figure 6F), with only WT female mice exhibiting kindling induced increased GFAP immunoreactivity (p=0.0157). G and H) There were no significant differences in NeuN immunoreactivity in CA1 and DG in female mice. I) Representative photomicrographs of immunohistochemical detection of GFAP and NeuN in mouse brain in WT and PSEN2 KO sham- and fully kindled mice. # Indicates significantly different from genotype-matched mice, p<0.05

In female mice (Figure 6E), both kindling (F_(1, 44)_ = 7.576, p=0.0086) and genotype (F_(1, 44)_ = 9.868, p=0.003) significantly impacted GFAP immunoreactivity in area CA1. *Post-hoc* analysis demonstrated stimulation-induced increases in GFAP immunoreactivity in WT female mice only (p=0.030). There was only a main effect of genotype on DG astrogliosis (F _(1, 45)_ = 7.057, p=0.0109; Figure 6F), with only WT female mice exhibiting kindling induced increased GFAP immunoreactivity (p=0.0157). As observed in male mice, there were no significant changes in NeuN immunoreactivity in either analyzed brain region.

## Discussion

This study provides critical demonstration that investigator-controlled induction of chronic focal seizures in an AD-associated preclinical model is suitable to interrogate functional and neuropathological impacts on disease trajectory, which may substantially benefit future disease modification strategies in patients with AD. This is the first conceptional application of the corneal kindling model of acquired epilepsy to directly assess the functional impacts of chronic seizures on AD-associated cognitive deficits. We have previously demonstrated that PSEN2 KO mice exhibit age-related changes in susceptibility to corneal kindling and pharmacological sensitively to antiseizure drugs [34]. We now extend our earlier findings to demonstrate the functional utility of kindling PSEN2 KO mice to assess the subsequent impacts of chronic seizures on cognitive decline (Table 2). The CKM is a moderate-throughput, minimally invasive model routinely used in epilepsy research [38, 52-56]. Unlike resource-intensive video-EEG recordings of spontaneous electrographic discharges with single-housed mice with AD-associated amyloid precursor protein (APP) overexpression [31, 57-63], corneal kindling induces network hyperactivity in an investigator-controlled setting wherein seizure history can be clearly identified; i.e. epileptogenesis can be directly controlled to time-dependently resolve how seizures affect disease outcomes at specific times throughout life [49]. Spontaneous seizures in epilepsy models are infrequent and unpredictable [64, 65], even with AD-associated mutations [63], therefore directly attributing behavioral changes to uniform seizure history is difficult, highly individualized, and requires large groups for long-duration, resource-intensive monitoring so as to have sufficient power to resolve an effect of seizures on behavior or disease modification [66, 67]. CKM can generate ample numbers of mice with uniform seizure history suited to cognitive testing [56] and pharmacological intervention [37, 41, 55]. This study provides the conceptual application to support such future studies in other AD-associated mouse models. Additionally, unlike long-term recordings of EEG-implanted rodents, CKM are group housed throughout their lifespan to minimize social isolation stress, which may itself induce behavioral deficits [68] and reduce seizure threshold of rodents [69]. Therefore, our present study substantially expands *in vivo* avenues to define the extent to which chronic seizures and AD-associated genotypes interact to modify cognitive function.

**Table 2.**
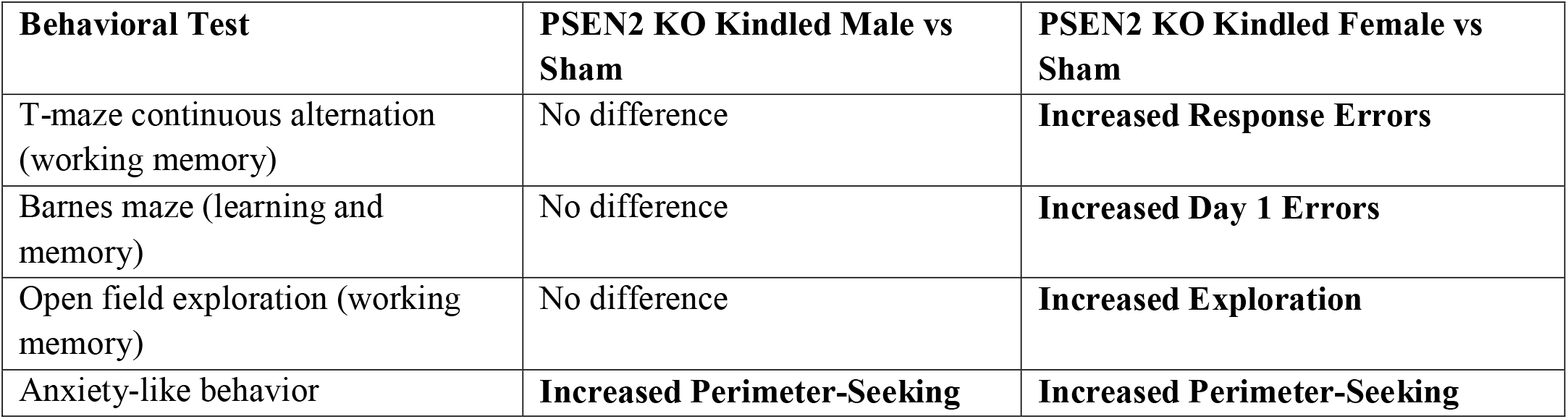
Male and female mice with loss of normal PSEN2 function (PSEN2 KO) aged over 6-months-old underwent corneal kindling or sham kindling to induce a history of chronic seizures before being challenged in several behavioral tasks to assess behavioral and functional impacts of chronic kindled seizures. Outcomes on each test are summarized below.

This work demonstrates that corneal kindled PSEN2 KO mice are useful to study functional impacts of chronic seizures in AD, as well as potentially uncover novel aging-related molecular drivers of epilepsy. We have previously demonstrated that young adult (2-3 months-old) PSEN2 KO mice are resistant to corneal kindling, but that this difference resolves by as early as 8-9-months-old [34]. We now report that loss of normal PSEN2 function may also prevent the formation of an epileptic network in mice aged 6-to 7-months-old. In alignment with our earlier work, seizure susceptibility of old (>14-months-old) PSEN2 KO female mice did not differ from age-matched WT controls, confirming that there are age-related changes in seizure susceptibility with loss of normal PSEN2 function in females. While we did not presently include aged (>14-months-old) male mice in this study nor assess functional impacts of kindled seizures in these old females, this work demonstrates that solely relying on younger-aged models with AD-associated genetic variants may miss critical and potentially important biological differences in seizure risk that arise with natural aging alone. Notably, reduced propensity for hyperexcitability has been previously demonstrated in an induced pluripotent stem cell model with loss of normal PSEN2 function [70]. However, despite resistance to kindling at this older age, evoked seizures were not more severe in PSEN2 KO mice. The kindled seizure was also stable at least 17 days after kindling acquisition during the period that all behavioral assays were performed. However, performance in several cognitively demanding, hippocampus-dependent tasks was only significantly worsened by evoked chronic seizures in aged female, but not male, PSEN2 KO mice. Our study adds to a growing body of literature to suggest that uncontrolled focal seizures could worsen disease burden in AD. Importantly, the cognitive burden of AD may be more substantially worsened by chronic, untreated focal seizures in women than in men.

PSEN2 KO mice are a unique platform on which to model the sequelae of chronic seizures in an AD-associated genotype [8] in the absence of accumulated Aβ. Prior studies of seizures in AD-associated rodent models have primarily relied on the pathological overexpression of APP to drive non-physiologic accumulation of Aβ [31, 59, 63]. PSEN2 KO mice are not defined by Aβ accumulation [71], this thus model provides a resource to define the Aβ-independent mechanisms underlying cognitive decline and advanced age in the setting of chronic seizures. While our study did not assess Aβ accumulation directly, subsequent studies are quantifying the impacts of chronic kindled seizures on Aβ accumulation in this and other AD-associated mouse models (*Vande Vyver, Barker-Haliski et al, submitted*). Importantly, PSEN2 variant models also allow for the specific interrogation of the role that inflammation may bidirectionally play on the cognitive and neuropsychiatric burden of AD, including functional outcomes from untreated focal seizures.

In this study, kindling-induced neuroinflammation was reproduced in 6-month-old WT male and female mice. However, chronic seizures only evoked hippocampal astrogliosis in male, but not female, PSEN2 KO mice. This study highlights major sex-specific impacts of uncontrolled chronic seizures on reactive astrogliosis. Considering that the most common PSEN2 variant (N141I) worsens inflammatory response of primary microglia [72], future work is needed to define whether chronic seizures and loss of normal PSEN2 function additively or synergistically modify other neuroinflammatory markers in a sex-specific manner. PSEN2 KO mice aged >6-months-old fill an important gap to define the extent to which chronic seizures and altered neuroinflammatory milieu can ultimately alter cognitive deficits with advanced age, as well as whether such impacts are sex-specific.

Hippocampal BDNF expression showed region- and sex-specific differences with loss of normal PSEN2 function and kindled seizures. BDNF is critical to neuroplasticity and may enhance α secretase and inhibit β secretase activities to effectively reduce Aβ production [73], and may reduce AD burden and delay disease pathology [74]. It is also disease-modifying in a rodent model of acquired epilepsy [75]. Given this neuroprotective and cognitive-sparing role of BDNF in both the AD-associated brain and epilepsy models, our present results demonstrating that BDNF expression in CA1 of the male hippocampus was not markedly upregulated in PSEN2 KO mice with chronic seizures starkly contrasted to the seizure-induced increased expression in age-matched WT mice (Figure 5A). Further, female mice did not show BDNF changes in CA1 in either genotype with kindling, suggesting sex-specific impacts on the expression of this neurotrophic factor with chronic seizures. There were also only significant reductions in BDNF expression in the DG of WT female mice, but no concomitant seizure-induced changes in PSEN2 KO mice. These data suggest that adult PSEN2 KO mice generally demonstrate disruptions in seizure-related changes in BDNF expression, with WT female mice generally exhibiting reduced hippocampal BDNF expression with kindling. Our findings of dysregulated BDNF expression in the PSEN2 KO model aligns other work demonstrating that the PSEN2-N141I variant is associated with reduced BDNF expression in an induced pluripotent stem cell model [70], as well as findings in an *in vivo* APP/PS1 model [76]. Dysregulated BDNF expression may thus contribute to the presently observed cognitive deficits exclusively detected in kindled female PSEN2 KO mice.

This present study highlights that loss of normal PSEN2 function may change propensity of epilepsy and/or hyperexcitability in an age-dependent manner and suggests that functional impacts can be independent of Aβ plaque formation. Our findings support a growing body of literature to suggest that APP and PSENs may modulate neuronal excitability through yet undefined mechanisms that are separate from their contributions to Aβ biogenesis [70]. Finally, PSEN2 variants are associated with later AD onset and may even be masked as sporadic AD cases [77], suggesting that PSEN2 variants could be more relevant to study late-onset AD than APP overexpression models. In this regard, our present study highlights the value of using an evoked CKM model to study seizure susceptibility and neurocognitive functional impacts across a lifespan. A significant percentage of patients with the most common AD-associated variant in PSEN2 also report seizures (32% of cases with the N141I variant) [13] and people with PSEN2 variants have high likelihood of seizures within 5 years of diagnosis [14], thus future studies must extend the present work to other PSEN-related AD variants to truly appreciate how uncontrolled chronic seizures affect functional and neuropathological outcomes in an age- and sex-related manner. This present study provides clear impetus to pursue the downstream consequences of seizures in the aging brain, and suggests that sex-specific therapeutic interventions to control focal seizures could be an opportunity to meaningfully slow cognitive decline in AD.

## Acknowledgements

This work was supported by the University of Washington Royalty Research Fund (MBH).

